# Machine Learned Classification of Ligand Intrinsic Activities at Human *µ*-Opioid Receptor

**DOI:** 10.1101/2024.04.07.588485

**Authors:** Myongin Oh, Maximilian Shen, Ruibin Liu, Lidiya Stavitskaya, Jana Shen

## Abstract

Opioids are small-molecule agonists of *µ*-opioid receptor (*µ*OR), while reversal agents such as naloxone are antagonists of *µ*OR. Here we developed machine learning (ML) models to classify the intrinsic activities of ligands at the human *µ*OR based on the SMILE strings and two-dimensional molecular descriptors. We first manually curated a database of 983 small molecules with measured *E*_max_ values at the human *µ*OR. Analysis of the chemical space allowed identification of dominant scaffolds and structurally similar agonists and antagonists. Decision tree models and directed message passing neural networks (MPNNs) were then trained to classify agonistic and antagonistic ligands. The hold-out test AUCs (areas under the receiver operator curves) of the extra-tree (ET) and MPNN models are 91.5 ± 3.9% and 91.8 ± 4.4%, respectively. To overcome the challenge of small dataset, a student-teacher learning method called tri-training with disagreement was tested using an unlabeled dataset comprised of 15,816 ligands of human, mouse, or rat *µ*OR, *κ*OR, or *δ*OR. We found that the tri-training scheme was able to increase the hold-out AUC of MPNN to as high as 95.7%. Our work demonstrates the feasibility of developing ML models to accurately predict the intrinsic activities of *µ*OR ligands, even with limited data. We envisage potential applications of these models in evaluating uncharacterized substances for public safety risks and discovering new therapeutic agents to counteract opioid overdoses.

**TOC Graphic:** 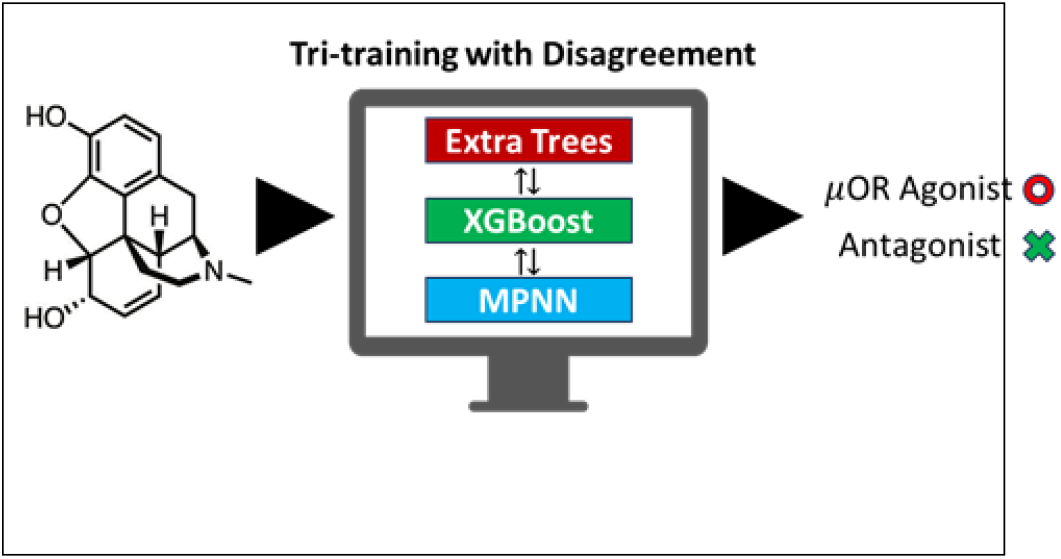

## Introduction

The opioid epidemic is a multifaceted and ongoing public health crisis that has resulted in countless deaths and significant societal and economic costs. ^1,2^ Despite various interventions, such as the prescription drug monitoring programs,^3^ medication-assisted treatment,^4^ naloxone distribution programs,^5^ and public education programs,^6^ opioid-related death rates have been sharply increasing year over year since 2019 (https://www.cdc.gov/opioids/basics/epidemic.html), highlighting the need for new and innovative approaches. By leveraging big data, machine learning (ML) models may help identify patterns, risk factors, and potential interventions. Recently, ML models have been applied to analyze a broad range of data sources, including electronic health records,^7^ social media data,^8^ and risk factors for opioid misuse^9,10^ and overdose. ^11^ Additionally, ML models ^12–14^ have been developed to analyze and predict the binding affinities of drugs to opioid receptors (ORs). Of particular interest is the study by Sakamuru et. al,^13^ in which several traditional ML models including tree models and support vector machine were built to predict small molecule agonists and antagonists of *µ*OR, *κ*OR, and *δ*OR. As training data, they measured the intrinsic activity (relative to DAMGO in agonist or naloxone in antagonist mode) of 2,805 compounds from the NCATS Pharmaceutical Collection of approved and investigational drugs^15^ using a quantitative high-throughput screening (qHTS) cAMP assay. However, it is unclear why antagonists and agonists were predicted separately, with the reported AUC of 88% for agonists and 76% for antagonists, while the balanced accuracy (BA) was 73% for agonists and 61% for antagonists.^13^

The present work aims to develop machine learning models for classification of agonistic and antagonistic activities of specifically the *µ*OR ligands. Different from the cAMP assay data used by the NCATS team, we manually curated a dataset of 983 human *µ*OR ligands with measured *E*_*max*_ values from the [^35^S]GTP*γ*S binding assay. The GTP*γ*S assay provides a more accurate measure of agonist activities compared to the cAMP assay, as it directly quantifies receptor occupancy at an early stage of the signaling cascade, avoiding potential amplification effects downstream of the receptor.^16^ Training on this dataset, the traditional tree models and directed message passing neural networks (MPNNs)^17^ were built for the classification task based on the SMILE representations of molecular structures. Graph neural networks (GNNs) are a class of neural networks that are designed to handle graph-structured data, and have shown great potential for various small molecule property prediction tasks in drug discovery, including solubility and toxicity prediction,^18–20^ protein-ligand binding affinity prediction,^21^ and antibiotic discovery.^22^ The directed MPNN uses a variant of the generic MPNN architecture tailored to use messages associated with directed edges (bonds) rather than messages associated with vertices (atoms).^17^ In 100 hold-out tests, our extra tree (ET) models and MPNNs achieved the average AUCs of 91.5% and 91.8%, respectively, and the average BAs of 83.3% and 85.1%, respectively. The MPNNs allowed us to interpret the model’s outcomes by identifying the smallest substructures responsible for antagonist activities. Finally, to overcome the barrier of small data size, we implemented a teaching-student learning model called tri-training with disagreement to improve model performance by leveraging a newly curated unlabeled dataset of 15,816 ligands of human, rat, and mouse ORs. Our work represents a beginning step of developing powerful ML models for predicting intrinsic activity at the OR.

## Results and Discussion

### Construction of the dataset

We compiled a comprehensive dataset of 983 human *µ*OR ligands from four online databases. We manually curated literature reports detailing the efficacy characterization of these compounds through [^35^S]GTP*γ*S functional assays of human *µ*OR. The author’s description was used to classify them as human *µ*OR agonist or antagonist. In cases where such descriptions were unavailable, we used naltrexone’s *E*_max_ value of 14% as a cutoff threshold for labeling the compounds (more details are given in Methods and Protocols). In total, we arrived at 755 agonists and 228 antagonist of human *µ*OR. The SMILE strings of these compounds were then used to calculate two-dimensional molecular descriptors using RDKit.^23^

### Analysis of the chemical space in the dataset

Using RDKit,^23^ we conducted a substructure analysis within the curated dataset and results are presented in Figure 1. The majority of these compounds are opioids and their core structures belong to six distinct groups (Figure 1): phenanthrene, benzomorphan, phenylpiperidine (PP), diphenylheptane, and phenylpropylamine, and others. The largest group of compounds (288) are phenanthrene derivatives, out of which 206 are agonists and 82 are antagonists. Many phenanthrenes are natural products found in the opium poppy plant. Morphine is a prototype opioid that belongs to this group, which contains other morphinans, 4,5*α*-epoxymorphinans, and oripavines. There are a total of 158 compounds (128 agonists and 30 antagonists) belonging to the benzomorphan group, which is obtained by removing the “C-ring” from morphine.^24^ Typical examples include ketocyclazocine, ethylketocyclazocine, phenazocine, and pentazocine. The majority of natural and semi-synthetic opioid drugs fall under the categories of phenanthrenes and benzomorphans. ^25^

**Figure 1:**
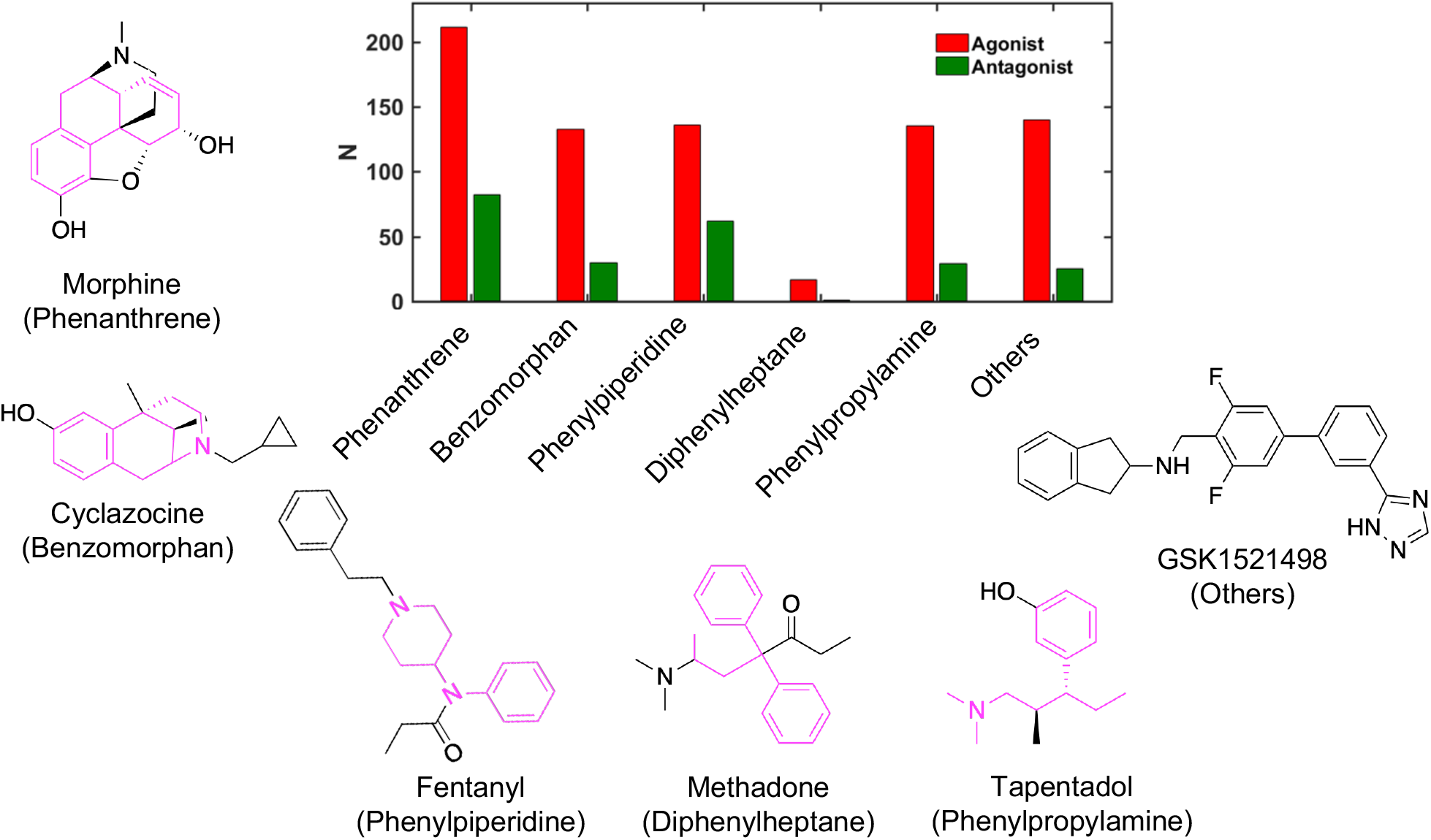
Opioid structure classes represented in the training dataset of human *µ*OR agonists and antagonists. The number (N) of agonists (red) and antagonists (green) that are derivatives of five major chemical classes, phenanthrene, benzomorphan, phenylpiperidine, diphenylheptane, and phenylpropylamine. All other molecules are grouped in Others. Commonly known opioids that are derivatives of these chemical classes (colored magenta) are shown. A novel synthetic opioid is shown as an example of Others.

In the newly constructed dataset, 132 agonists and 62 antagonists belong to the phenylpiperidine opioid class. Most synthetic opioids are often less complex molecules without multiple rings in the backbone. One of the highly potent types of synthetic opioids are phenylpiperidine derivatives, ^26^ including fentanyl, alfentanil, and sufentanil. 4-anilidopiperidines and their analogues are also included in the phenylpiperidine group. The diphenylheptane group contains the least number of compounds, only 17 agonists and 9 antagonists. Examples of chemicals found in this class are methadone and propoxyphene. The tramadol group contains 132 agonists and 29 antagonists, and includes tramadol and tapentadol. A sizable number of compounds (137 agonists and 25 antagonists) do not belong to any of the aforementioned classes.

To further understand the chemical space of the training dataset, we performed a clustering analysis of the chemical structures based on the Tanimoto distances (1-Tanimoto similarity index)^27,28^ between two molecules’ SMILES strings. A hierarchical agglomerative method in Scikit-learn^29^ was used, with a Tanimoto distance of 0.7 as the clustering cutoff. A total of 23 clusters were found; the relative populations and the corresponding percentages of antagonists in the clusters are shown in Figure 2. The top (largest) two clusters (Cluster #1 and #2) contain about 35% and 20% of the total number of molecules in the dataset, whereas the rest of the clusters each contain less than 10% of the dataset. About 30% of the molecules in the top two clusters are antagonists, which reflects the 1:3.3 antagonist:agonist ratio of the dataset (Figure 2). The clustering analysis also allowed us to identify antagonists that are structurally similar to the agonists. For example, the most representative molecule (i.e., cluster centroid) of cluster #1, asalhydromorphone is a prodrug of a potent narcotic hydromorphone. Asalhydromorphone is structurally similar to naldemedine tosylate, which is a *µ*OR antagonist used for treatment of opioid-related constipation ^30^ (Figure 2, cluster #1). The percentages of antagonists allowed us to identify a cluster dominated by antagonists, i.e., cluster #11 which contains 13 molecules with 10 (77%) being antagonists. The representative molecule of cluster #11 is an antagonist; its structure and the most similar structure of an agonist are displayed in Figure 2.

**Figure 2:**
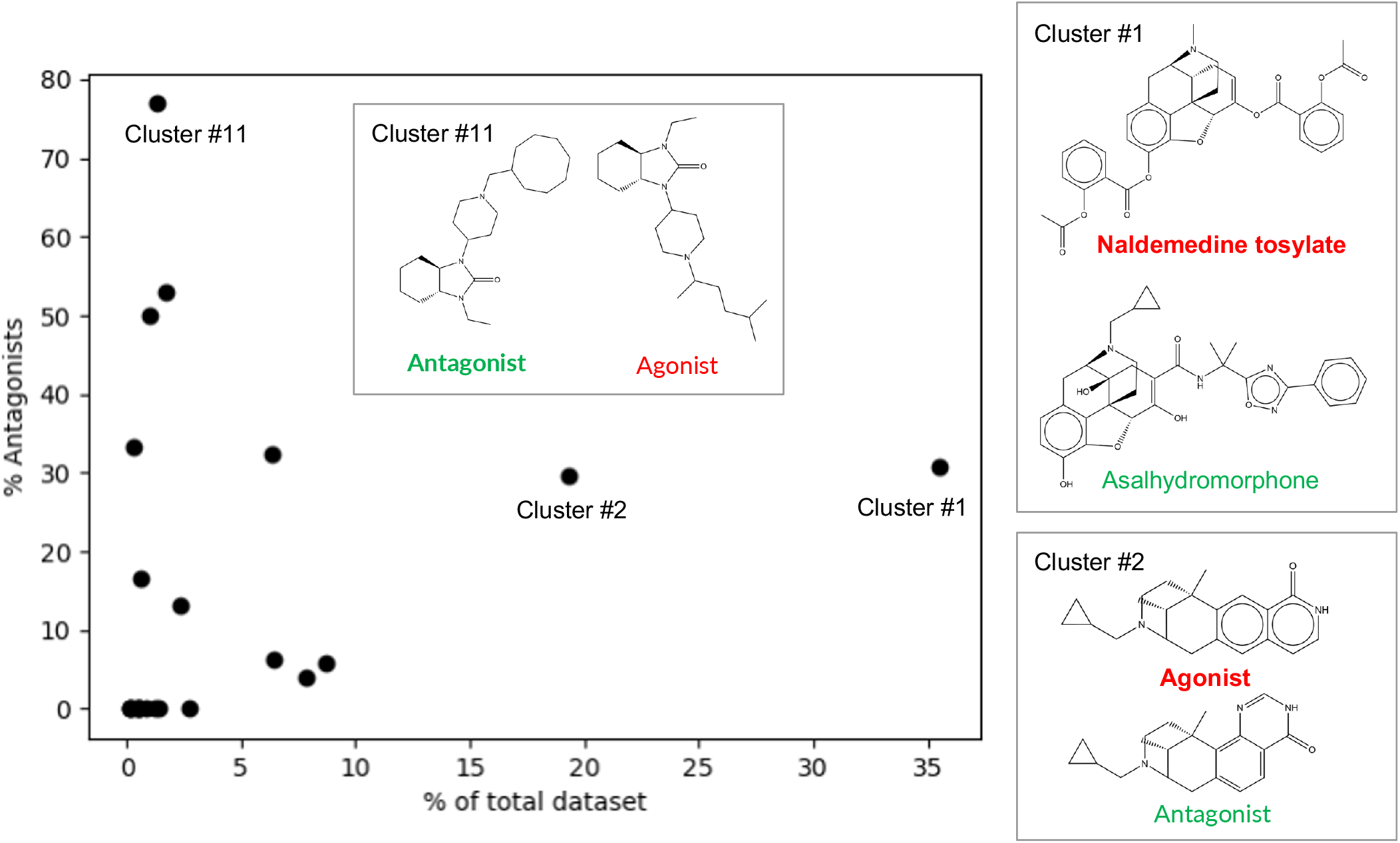
Clustering analysis of the chemical structures in the training dataset of human *µ*OR agonists and antagonists. Relative population of the cluster vs. the percentage of antagonists. The clusters are numbered based on the populations. For Cluster #1, #2, and #11, the representative structure (cluster centroid, bold font) and the most similar structure (highest Tanimoto similarity) of a molecule from the opposite functional class are shown. The named molecules are discussed in the main text.

### Performance of the decision tree models

PyCaret^31^ was used to train the random forest (RF) and extra tree (ET) models for binary classification of a compound as agonist or antagonist. Since the number of agonists is 3.3 higher than that of the antagonists, the latter were labeled as positives and agonists were labeled as negatives in the model training. The dataset was randomly split in a 9:1 ratio for training/cross-validation (CV), and unseen hold-out testing. Each molecule was represented by the SMILE string and the two-dimensional molecular descriptors calculated from RDKit.^23^ To address the data imbalance issue, a minority oversampling technique SMOTE^32^ was employed to generate synthetic data points for the minority class (antagonists). After removing the highly correlated and invariant descriptors, the feature space was reduced to 91. To further reduce model overfitting, we built and tested the models with 9 smaller feature sets using the feature selection threshold *α* between 0.2 and 1 in PyCaret, ^31^ which corresponds to 49–91 features. The data splitting, model training and evaluation were performed 100 times to ensure a robust statistical analysis of the model performance. We examined the area under the curve of receiver operator curve (AUC-ROC or AUC), balanced accuracy (BA), and F1 score (for the minority class) for the resulting models (Supplemental Table S1).

The performances demonstrated minimum dependence on the size of feature space. The AUCs range 91.0–91.8% and 90.3–91.5%, the BAs range 81.7-83.1% and 80.7–83.3%, and the F1s range 72.6–74.9% and 71.3–75.4% for the RF and ET models, respectively (Supplemental Table S1). The highest AUC of 91.8 ± 2.7% for the RF and 91.5 ± 3.9% for the ET model was achieved with 90 features (*α* = 0.8). The highest BA of 83.1 ± 5.2% was obtained for the RF model with 70 features (*α* = 0.4) and 83.3 ± 5.0% for the ET model with 90 features (*α* = 0.8). In terms of F1, the models attained their highest values of 74.9 ± 6.4% for the RF model with 49 features (*α* = 0.2) and 75.4 ± 6.4% for the ET model with 90 features (*α* = 0.8).

To further evaluate the RF and ET models with the highest BA, we examine the recall and precision for both agonist and antagonist classes (Table 1). The recall and precision of predicting agonists (the majority class) are very high, 93.3 ± 2.8% and 92.0 ± 2.7% for the RF model, and 94.1 ± 3.0% and 92.0 ± 2.6% for the ET model. The recall and precision of predicting antagonists (the minority class) are lower, 72.9 ± 9.9% and 77.0 ± 8.1% for the RF model, and 72.6 ± 9.9% and 79.5 ± 8.1% for the ET model. Given the over 3:1 ratio for agonists:antagonists, this difference in performance is expected (despite the use of the minority oversampling technique). Nonetheless, the average AUCs of the RF and ET models are over 91%, suggesting that the models have high classification power. In the work of Sakamuru et al.,^13^ a majority undersampling scheme was used and the RF model gave the highest performance metrics among all the classifiers tested, including support vector machine, (simple) neural network, extreme gradient boosting and RF. Specifically, the model for the *µ*OR agonists achieved the highest AUC of 88% and BA of 73%.^13^ Despite a smaller training dataset, our RF model has a higher performance, with the average AUC of 91% and BA of 83.1% (Table 1).

**Table 1:** Summary of the evaluation metrics for the best RF and ET models on the hold-out data.

|  | RF (%) | ET (%) |
| --- | --- | --- |
| Recall(AG) | $93.3 \pm 2.8$ | $94.1 \pm 3.0$ |
| Recall(AN) | $72.9 \pm 9.9$ | $72.6 \pm 9.9$ |
| Prec(AG) | $92.0 \pm 2.7$ | $92.0 \pm 2.6$ |
| Prec(AN) | $77.0 \pm 8.1$ | $79.5 \pm 8.1$ |
| BA | $83.1 \pm 5.2$ | $83.3 \pm 5.0$ |
| F1(AN) | $74.5 \pm 7.4$ | $75.4 \pm 7.2$ |
| AUC | $91.0 \pm 3.8$ | $91.5 \pm 3.9$ |

### Analysis of important functional groups for model predictions

To rationalize the predictions made by the tree models, we computed SHAP (SHapley Additive exPlanations) values, ^33^ which employ a game theoretical approach to quantify the impact of each feature to the model’s output. Figure 3 displays the top 10 features ranked by their mean absolute SHAP values on the vertical axis, while the horizontal axis depicts the SHAP values of each feature for a set of instances in the dataset. The SHAP values represent the contribution of each feature to the difference between the model’s predicted output and the average output. Positive SHAP values signify a change in the expected model prediction towards antagonists.

**Figure 3:**
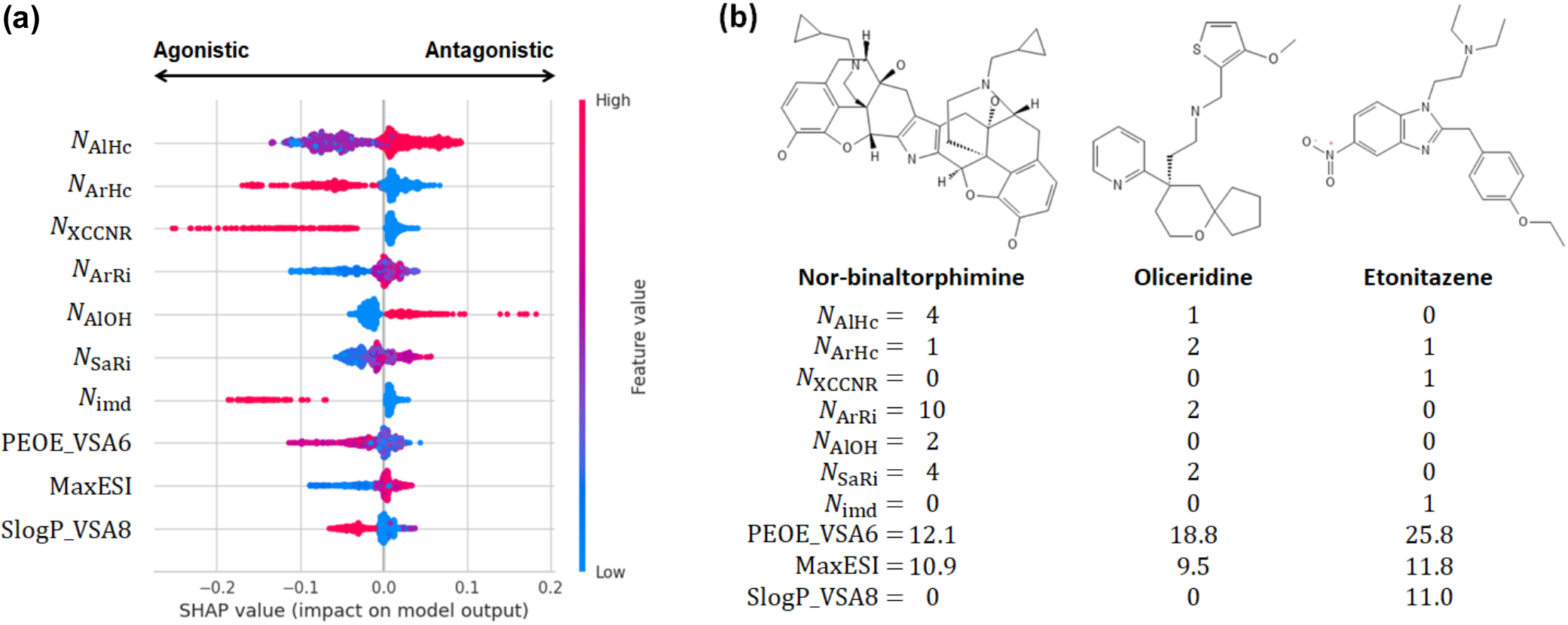
Analysis of important features of the ET model. **(a)** SHAP beeswarm plot for the 10 highest ranked features in the descending order of the mean absolute SHAP values for the best performing ET model (Table 1). Each dot corresponds to one instance in the dataset. The color scale represents the magnitude of the feature. The top 7 features are the number of aliphatic heterocycles, aromatic heterocycles, XCCNR groups, aromatic rings, aliphatic OH groups, saturated rings, and imidazole groups. The bottom 3 features are the charge descriptor, maximum electrotopological state index, and the logP descriptor. Detailed explanation of the features are given in Table S2. **(b)** Molecular structures of nor-binaltorphimine (antagonist), oliceridine (agonist), and etonitazene (agonist) and their feature values.

Based on the average SHAP values, the seven most important features are the number of aliphatic (*N*_AlHc_) and aromatic heterocycles (*N*_ArHc_), XCCNR groups (tertiary amine, *N*_XCCNR_), aromatic rings (*N*_ArRi_), aliphatic OH groups (*N*_AlOH_), saturated rings (*N*_SaRi_), and imidazole groups (*N*_imd_). The SHAP beeswarm plot reveals that the number of aliphatic OH groups makes the largest positive impact on the model output, and specifically, larger number of OH groups shifts the model prediction towards antagonists (Figure 3a). The second largest positive impact is made by the number of aliphatic heterocycles; a larger number of aliphatic heterocycles also shifts the model prediction towards antagonists (Figure 3a). As to the features that make large negative impacts on the model output, the SHAP beeswarm plot reveals that the number of XCCNR groups makes the largest negative impact followed by the number of aromatic heterocycles and the number of imidazole groups (Figure 3a). Increasing values of these features shift the model predictions towards agonists (Figure 3a).

To corroborate the SHAP value analysis, we examine the distributions of the feature values for the antagonists and agonists (Figure S2) and three labeled compounds in the training set (Figure 3b). *N*_AlHc_ has modes 0, 1, and 2, with mode 1 and 2 dominated by the antagonists and mode 0 dominated by the agonists (Supplemental Figure S2). This is consistent with large *N*_AlHc_ values shifting the predicted outcomes to antagonists (Figure 3a). For example, the antagonist nor-binaltorphimine has the largest *N*_AlHc_ value of 4, as compared to 1 for oliceridine and 0 for etonitazene, both of which are agonists (Figure 3b). *N*_AlOH_ has modes 0 to 4, with mode 0 and 1 dominated by the agonists and mode 2 and 3 dominated by the antagonists (Supplemental Figure S2). These distributions are consistent with larger *N*_AlOH_ values predicting antagonists (Figure 3a), e.g., in the three examples, the only antagonist norbinaltorphimine has *N*_AlOH_ of 2, whereas the agonists oliceridine and etonitazene have *N*_AlOH_ value of 0 (Figure 3b). Similarly, we found that the modes of *N*_XCCNR_, *N*_ArHc_, and *N*_imd_ (Supplemental Figure S2) corroborate the SHAP beeswarm plot as well as the labels of the three example compounds, demonstrating that increased values shift the predicted outcomes to agonists (Figure 3a).

### Performance of the MPNN models

GNNs are expected to be superior than the traditional tree models for large datasets. To test this hypothesis, we trained the MPNN models based on the molecular graphs built from the SMILE strings and the default settings in ChemProp. ^34^ The performance of these initial MPNN models is similar to the ET models, with the average BA of 83.1%, F1(AN) of 75.4%, and AUC of 90.8 % (Table 2). Next, we tested several strategies to improve the model performance: adding molecular features from RDKit,^23^ optimizing the classification threshold, and tuning the hyperparameters (Table S3).

**Table 2:** Summary of the evaluation metrics for the initial and optimized MPNNs on the holdout data.

| Model | Initial | Optimized |
| --- | --- | --- |
| Recall(AG) | 94.5 ± 2.5 | 93.9 ± 3.1 |
| Recall(AN) | 71.7 ± 10.0 | 76.3 ± 10.6 |
| Prec(AG) | 91.7 ± 2.7 | 93.0 ± 2.9 |
| Prec(AN) | 80.2 ± 7.7 | 79.5 ± 8.1 |
| F1(AN) | 75.4 ± 7.3 | 77.5 ± 6.7 |
| BA | 83.1 ± 5.1 | 85.1 ± 5.0 |
| AUC | 90.8 ± 3.8 | 91.8 ± 4.4 |

The average and standard deviations of the metrics from 50 runs of data splitting and model training/CV are given in percentages.

Compared to the initial MPNN, classification threshold optimization improved the recall of antagonists but decreased the F1 (Model 1 and 2, Table S3 and S4). However, adding molecular features increased the recall by 0.5% and 0.7% and the precision by 1.2% and 0.3% for the antagonist and agonist classes, respectively (Model 1 and 3, Table S3 and S4). As a result, the F1 score was increased by 1.0% (Table S4). Next, the classification threshold was varied to maximize the F1 score, which led to a 0.2% increase of F1 and 0.8% increase of the BA (Model 3 and 4, Table S4). This increase was driven mainly by the significant (3.1%) increase in the recall of antagonists and to a smaller extent the precision improvement for predicting the agonists (0.6% increase); however, the recall of agonists and precision for antagonists were decreased by 0.6% and 2.9%, respectively (Model 3 and 4, Table S4). While the above metrics were obtained using ensemble training of 10 models from the 10-fold CV, we also tested ensemble training of 5 models (5-fold CV), which resulted in only a small decrease of 0.2% for the F1 score (Model 4 and 5, Table S4). Consequently, to reduce model training time, we proceeded to hyperparameter optimization using the ensemble training of 5 models. This step gave a significant boost in the model predictions for both classes, with the F1 and BA increasing by 1.1% and 1.8%, respectively (Model 5 and 6, Table S4). The resulting optimized MPNN (model 6 in Table S4) shows significantly higher performance than the best tree model (ET), with the F1 of 77.5 ± 6.7% and BA of 85.1 ± 5.0%, as compared to the ET’s F1 of 75.4 ± 7.2% and BA of 83.3 ± 5.0%. The AUC of the optimized MPNN is 91.8 ± 4.4%, which is slightly (0.3%) higher than the ET’s AUC of 91.5 ± 3.9%.

### External validation and analysis of predictions

To further test the predictive power of the models, we refined the ET (the best tree model) and MPNN by training on the entire dataset of 755 agonist and 228 antagonists of *µ*OR and applied the final models to predict on an external set of 11 *µ*OR antagonists and 16 agonists that were recently published but not in the model training set (Supplemental Table S5). Overall, the MPNN outperformed the ET model in all metrics, consistent with the hold-out test results. The ET model correctly predicted 11 out of 15 agonists and 7 out of 11 antagonists, giving the respective recalls of 73.3% and 63.6% with a balanced accuracy of 68.5% (Table 3). The MPNN correctly predicted 12 out of 15 agonists and 9 out of 11 antagonists, giving the respective recalls of 80.0% and 81.8% with a balanced accuracy of 80.9% (Table 3). Thus, the MPNN model has a higher recall for both classes than the ET model, particularly the minority antagonist class, with an increase of 18.2%. The precision of the MPNN model is also significantly higher than the ET model, particularly the minority antagonist class, with an increase of 11.4%.

**Table 3:**
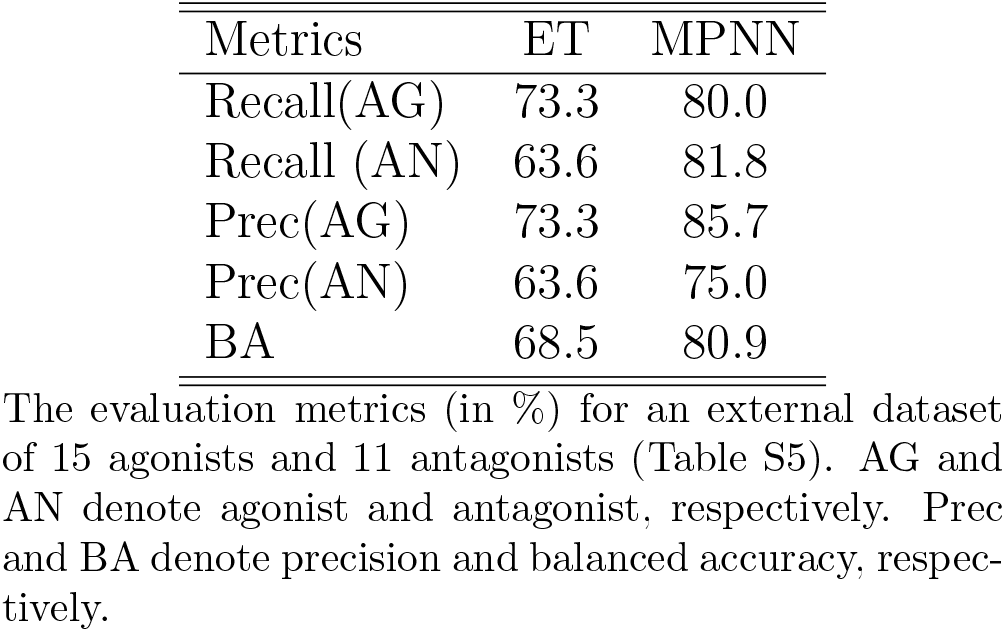
Performance of the final ET and MPNN models on the external dataset of 26 compounds.

To rationalize the predictions, we analyzed the structures of the external compounds and applied the Monte-Carlo tree search algorithm implemented in ChemProp^34^ to identify substructures responsible for the antagonistic classifications. The external dataset contains 5 phenanthrene-based antagonists, 6*β*-naltrexol, nalodeine (*N*-allylnorcodeine), (17S)-methylnaltrexone, naloxazone, and orvinol 14. Figure 4 shows the substructures responsible for the antagonist predictions. The MPNN correctly classified 4 out of 5 phenanthrene antagonists. The model mis-classified orvinol 14 as agonist, perhaps based on the high similarity with the structure of BU08028 (Tanimoto score of 0.99), which is an agonist in the training set (Figure 4).

**Figure 4:**
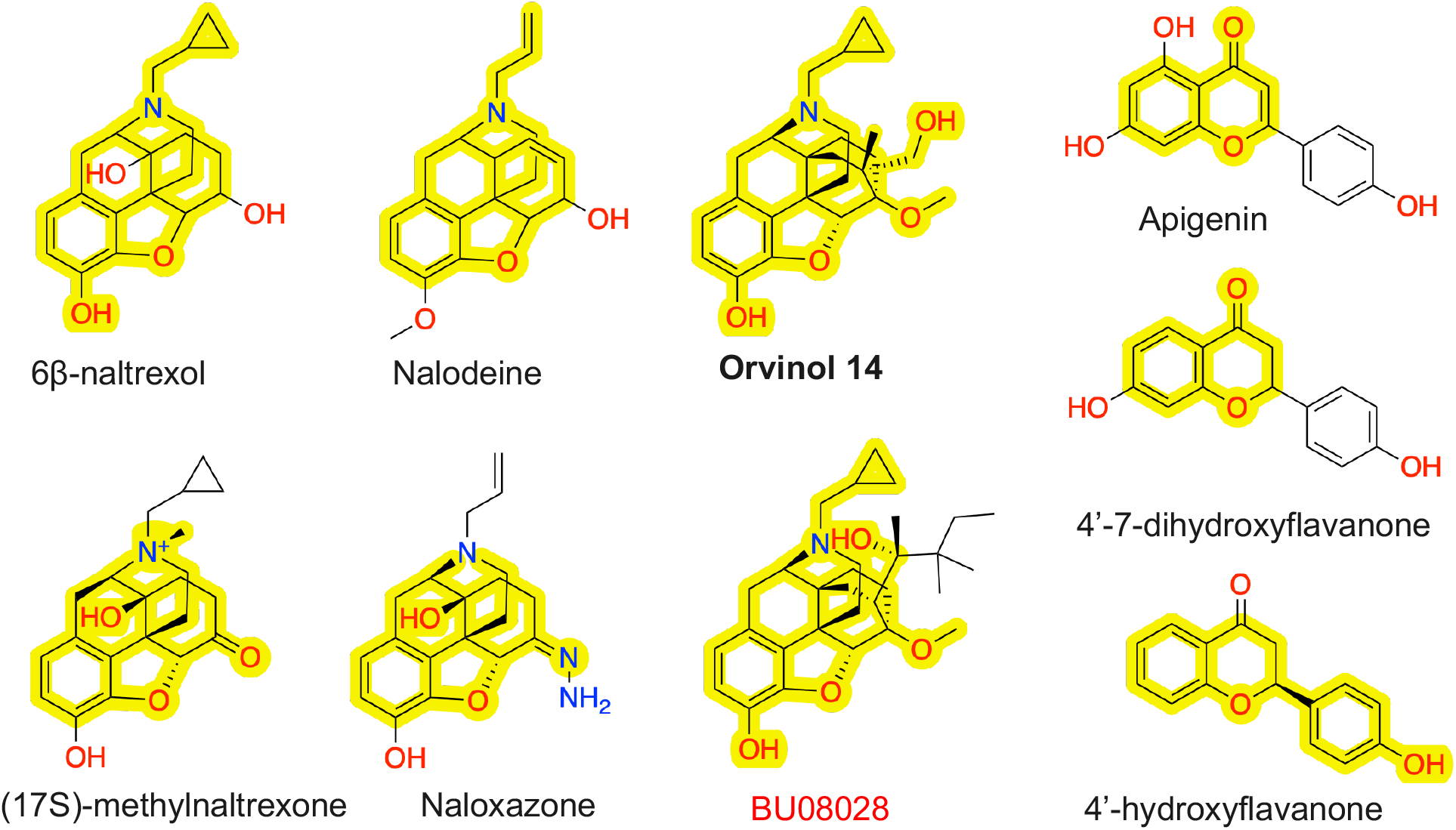
Chemical structures of the phenanthrene-based (a) and flavonoid (b) antagonists in the external validation set. The key substructures responsible for the model prediction of antagonist are highlighted. Molecules in bold are incorrectly predicted as agonist. The key substructure of orvinol 14 is found in the agonist BU08028 (red) in the training set (common substructure is highlighted).

The external dataset also contains three flavonoids, apigenin (or 4’,5,7-trihydroxyflavone), 4’,7-dihydroxyflavone, and 4’-hydroxyflavanone (Figure 4). Flavonoids represent an important class of natural products that belong to plant secondary metabolites with a polyphenolic structure. ^35^ The Monte-Carlo tree search algorithm pinpointed chromone (1,4-benzopyrone), which is a part of the 2-phenylchromen-4-one backbone of flavones, as the key substructure for the antagonistic predictions. The MPNN correctly classified apigenin and 4’,7-dihydroxyflavone, which differ by a hydroxyl group, as antagonist. However, the MPNN model classified 4’-hydroxyflavanone as agonist (Figure 4).

The external validation dataset contains four phenanthrene-based agonists, 3-monoacetylmorphine, morphinone, codeinone, and buprenorphine hemiadipate, which were all correctly classified by the MPNN, while the ET model misclassified one agonist, buprenorphine hemiadipate (Figure 5). To interpret the model predictions, we note that the four phenanthrene-based agonists are derivatives of morphine (Figure 5), which is in the training set. Moreover, the methyl or acetyl substitution at the C3 position in 3-monoacetylmorphine, codeinone, and buprenorphine hemiadipate is seen in methylmorphine (codeine), ethylmorphine, and diacetylmorphine (also known as heroin), while buprenorphine hemiadipate is a C3-substituted buprenorphine (Figure S3). It is noteworthy that both the MPNN and ET models incorrectly classified three agonistic compounds, morphiceptin, matrine and sophocarpidine (Figure 5). Morphiceptin is a peptide, while matrine and sophocarpidine are alkaloids. We suggest that the erroneous predictions may be related to the lack of structurally similar agonistic compounds in the training dataset.

**Figure 5:**
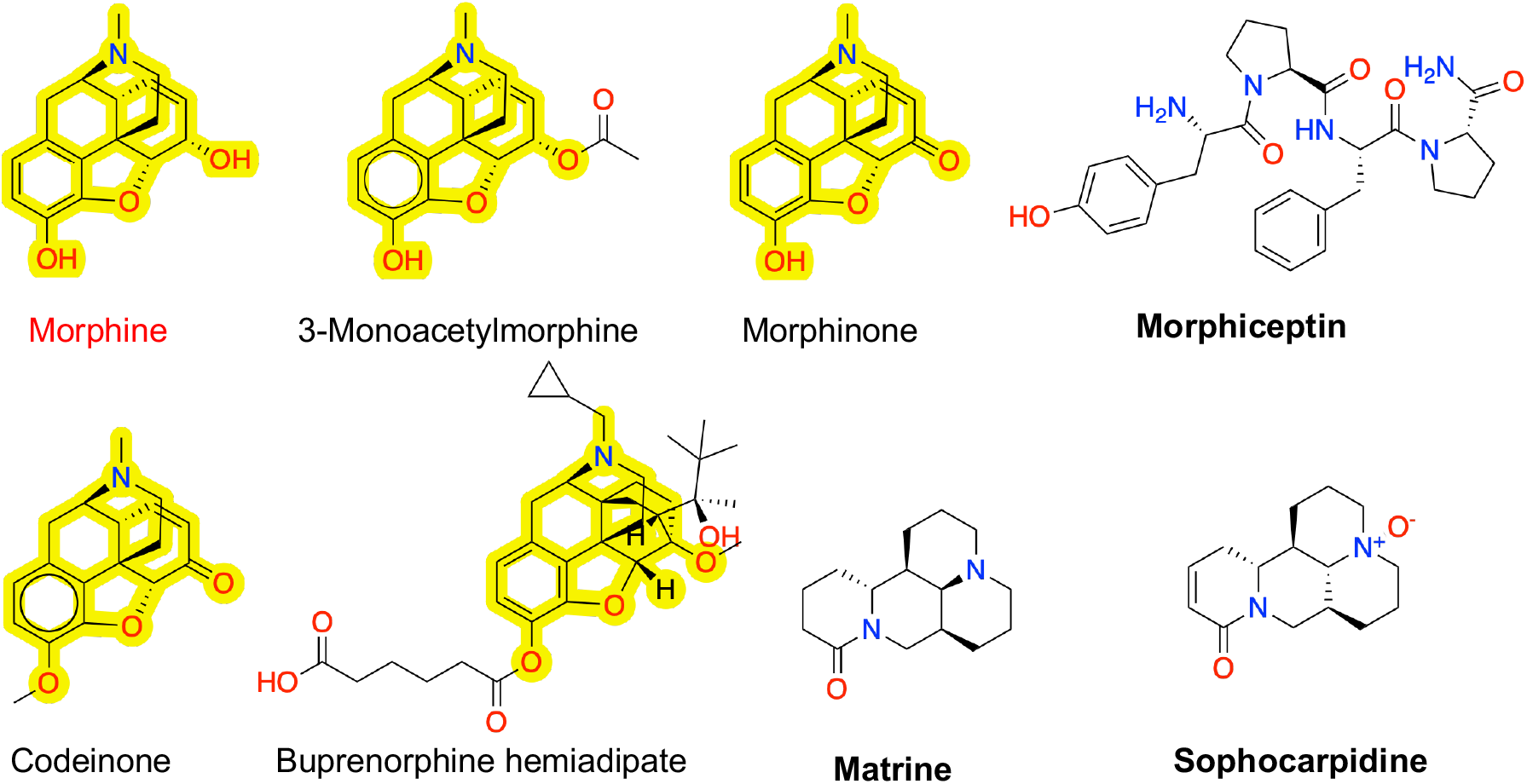
Molecular structures of the phenanthrene-based agonists and three incorrectly classified agonists in the external validation dataset. The phenanthrene-based agonists are derivatives of morphine (red). The morphine substructure is highlighted in these compounds. The three mis-classified agonists are in bold font.

### Proof of concept of a tri-training scheme with unlabeled data to improve models

To overcome the issue of small data size and improve the model performance, we implemented a semi-supervised teacher-student learning scheme called tri-training with disagreement.^36,37^ We first curated an unlabeled dataset comprised of 15,816 ligands of human, mouse, or rat *µ*OR, *κ*OR, or *δ*OR taken from bindingDB,^38^ which is 15 times larger than the labeled training dataset comprised of 983 ligands. A tri-training run began by stratified random splitting of the labeled data to 90% training/CV and 10% hold-out test. Next, three different models, ET, XGBoost, and MPNN were used to classify an unlabeled ligand. If one model disagreed with the other two, its original (labeled) training set was supplemented by the ligand and the pseudo label given by the other two models. If all three models agreed, the ligand was discarded. This process continued until the unlabeled dataset was depleted, at which point all three models were retrained and cross-validated using the supplemented training set. This was then followed by testing on the 10% labeled hold-out data. In the next iteration, the new models were used to classify the discarded data, and the iterations continued until the AUC no longer improved, unlabeled dataset was depleted, or the three models no longer disagreed.

As a proof of concept, we performed three tri-training runs with different training/CV-testing data splits (9:1 ratio) and no hyperparameter tuning. Following the hold-out tests after each iteration, the models were externally validated using the aforementioned 26 compounds (Table S6). In Run 1, the three initial XGB, ET, and MPNN models have the hold-out test AUCs of 90.6%, 86.4%, and 89.7%, respectively (Table 4). After the first iteration, the hold-out test AUCs of the XGB and ET models significantly increased to 94.2% and 93.1%, respectively (Table 4). In the subsequent iterations, the AUCs of both models oscillated between decreasing and increasing. After iteration 5, the hold-out test AUCs of the XGB and ET models were 94.9% and 92.3%, respectively, which are substantially higher than the respective initial model AUCs (Table 4). However, only the external validation AUCs of the XGB models were higher than the initial model AUC after tri-training iterations, with the highest AUC found after the first iteration at 87.9%, as compared to 69.6% for the initial model (Table S6). In contrast, the external validation AUCs of the ET model after tri-training iterations were all lower than the initial model AUC.

**Table 4:** Summary of the AUCs of the holdout tests from three runs of tri-training with disagreement.

| Iter | Data | XGB | Data | ET | Data | MPNN |
| --- | --- | --- | --- | --- | --- | --- |
| Run 1 |  |  |  |  |  |  |
| 0 | 886 | 90.6 | 886 | 86.4 | 886 | 89.7 |
| 1 | 1683 | 94.2 | 1673 | 93.1 | 5416 | 88.8 |
| 2 | 2287 | 89.5 | 2834 | 90.1 | 6173 | 89.4 |
| 3 | 2417 | 91.9 | 2861 | 91.4 | 6351 | 88.4 |
| 4 | 2455 | 91.5 | 2865 | 91.9 | 6489 | 90.4 |
| 5 | 2477 | 94.9 | 2872 | 92.3 | 7005 | 89.8 |
| Run 2 |  |  |  |  |  |  |
| 0 | 886 | 84.4 | 886 | 90.5 | 886 | 92.3 |
| 1 | 1755 | 92.9 | 1727 | 90.1 | 6436 | 92.6 |
| 2 | 2977 | 80.7 | 2312 | 84.5 | 6796 | 90.6 |
| 3 | 2991 | 78.8 | 2344 | 84.6 | 6883 | 92.8 |
| 4 | 2995 | 88.6 | 2363 | 83.5 | 6886 | 93.1 |
| 5 | 2996 | 85.0 | 2373 | 82.7 | 6894 | 93.7 |
| Run 3 |  |  |  |  |  |  |
| 0 | 886 | 86.3 | 886 | 82.3 | 886 | 93.3 |
| 1 | 2263 | 87.0 | 1705 | 85.1 | 4592 | 94.5 |
| 2 | 2681 | 93.7 | 3387 | 85.3 | 4979 | 95.4 |
| 3 | 2737 | 94.4 | 3396 | 84.6 | 5181 | 95.2 |
| 4 | 2771 | 88.4 | 3397 | 84.0 | 5283 | 95.7 |
| 5 | 2784 | 93.5 | 3398 | 83.4 | 5391 | 95.1 |

Interestingly, the tri-training results of MPNN showed a very different trend compared to those of the tree models. The test AUC only increased slightly after iteration 4. This difference between the MPNN and tree models is consistent with the different training data size at each iteration. The training dataset size for each iteration converged rapidly, with the XGB and ET models receiving 22 and 7 additional data points, respectively in iteration 5 (Table 4). However, the large amount of data points being added to the MPNN training set after the first iteration (Table 4) suggests that the XGB and ET models were initially too similar, and agreed very often whereas MPNN disagreed.

In the tri-training run 2, the initial AUCs for the XGB, ET, and MPNN models are 84.4%, 90.5%, and 92.3%, respectively. The behavior of data size expansion is very similar to that of run 1, supporting our conclusion that the two tree models are similar, whereas MPNN is different. Interestingly, both the test and external validation AUCs of the XGB model significantly increased after iteration 1, whereas the test and external validation AUC of the ET model did not increase in any iteration (Table 4 and Table S6).

As to the MPNN model, the changes in the test AUC were smaller compared to the tree models; however, except for iteration 2, the test AUCs after tri-training were all higher than the initial value. After the first iteration, the test AUC was 92.6%, as compared to 92.3% of the initial model (Table 4). The corresponding external validation AUC was 84.2%, as compared to 83.0% of the initial model (Table S6).

In the tri-training run 3, the initial AUCs for the XGB, ET, and MPNN models are 86.3%, 82.3%, and 93.3%, respectively. The behavior of data size expansion was still very similar to the previous two runs, with slightly less data being added to the MPNN training set. The test AUCs of all three models significantly increased with iterations, to maximums of 94.4%, 85.3%, and 95.7% respectively. However, the external validation AUCs did not follow the same trend, and only saw significant improvement after iteration 1 of the XGB model. The inconsistency between the hold-out test and external validation performances is likely due to the extremely small size of the latter. Nonetheless, these data suggest that tri-training with disagreement is a promising strategy for increasing the model performance given a small labeled dataset.

## Concluding Discussion

We developed ML models to classify the intrinsic activities of small molecule ligands of the human *µ*OR by training on a manually curated dataset comprised of 983 *µ*OR ligands including 755 agonists and 228 antagonists that have literature *E*_max_ values using the [^35^S]GTP*γ*S functional assay. The ET and MPNN models demonstrated high predictive performance in 100 unseen tests, achieving AUCs of 91.5 ± 3.9% and 91.8 ± 4.4%, respectively, indicating their capability to accurately discriminate between agonists and antagonists, with MPNN being slightly more superior. The recall/precision for agonists are 94.1 ± 3.0%/92.0 ± 2.6% with ET and 93.9 ± 3.1%/93.0 ± 2.9% with MPNN, while those for antagonists are 72.6 ± 9.9%/79.5 ± 8.1% with the ET and 76.3 ± 10.6%/79.5 ± 8.1% with the MPNN. The decrease in the prediction performance for antagonists is due to the significant (3:1) class imbalance. The finalized ET and MPNN models, trained on the entire dataset, achieved BAs of 68.5% and 80.9%, respectively, on an external validation set of 26 *µ*OR ligands (15 agonists, 11 antagonists). The MPNN significantly outperformed the ET, with over 80% recall for both agonists and antagonists.

Aiming to overcome the limitations of class imbalance and small training dataset, we tested a teacher-student learning method, tri-training with disagreement, to enhance model performance by using the unlabeled data comprised of ligands of *µ*OR, *κ*OR, and *δ*OR from human, mouse, and rat. In all three runs, the model test and validation AUCs in some runs were increased, demonstrating the potential of the tri-training scheme. In particular, the test and external validation AUCs of the XGB model were substantially increased after just one tri-training iteration, where the training data size nearly doubled through receiving pseudo-labeled data predicted by the other two models. The tri-training data also suggests that the two tree models are very similar whereas the MPNN differs from the two, which resulted in a dramatic expansion of the training data size of the MNPP model after just one iteration (addition of about 4500 pseudo-labeled data points as compared to about 800 for the two tree models). Judging by the change in the test and validation AUCs, the tri-training effect on the MPNN is minimum, which can be attributed to the lower performance of the initial tree models. By the same token, the learning outcome of the XGB model is most obvious (run 3), which can be attributed to the higher quality of teachers, i.e., the ET and MPNN giving out accurate labels.

The current models are limited to predicting actives, as the training data exclusively comprised experimentally validated human *µ*OR ligands. To address this limitation and improve performance, future efforts may be directed at expanding the training dataset with diverse, experimentally confirmed *µ*OR-inactive compounds and reformulating the ML task as a multiclass classification problem, training the model to distinguish between three pharmacologically distinct categories: inactives, agonists, and antagonists of the human *µ*OR. Another limitation of the models is that only binders are considered. We envision a two-step protocol addressing binder/non-binder prediction followed by pharmacological effect prediction. For binder/non-binder classification, molecular docking and the use of a decoy set could complement ML models.

Labeling compounds as agonists or antagonists based on an empirical threshold (e.g., 14% *E*_max_ value) or literature conclusions oversimplifies the complexity of ligand pharmacological profiles,^39^ neglecting the nuanced range from inverse agonism to full agonism, e.g., the subtle differences between weak partial agonists and antagonists. Although regression models can in principle be developed to predict *E*_max_ values, such effort is impeded by the variability in the reported measurements across different published studies. This variability arises from various factors, including differences in receptor expression levels, the specific cell lines used, and experimental conditions employed by different research groups.

Despite the aforementioned limitations, over 91% AUC values for both the ET and MNPP models indicate that they are capable of accurately classifying the intrinsic activities of small molecule ligands of *µ*OR. Such models may find many important applications, for example, help discover new *µ*OR antagonists for treatment of opioid overdose. Additionally, they may be used to evaluate pharmacologically uncharacterized substances that may pose a risk to public safety.

## Methods and Protocols

### Training, testing, and external validation datasets

We first identified four online databases that contain opioid structures, including ChEMBL,^40^ GLASS,^41^ BindingDB, ^42^ and IUPHAR/BPS Guide to Pharmacology ^43^ We then collected a total of 983 chemical compounds that interact with exclusively human *µ*OR from these databases. We labeled a compound as agonist or antagonist based on the authors’ description in the cited literature reference, resulting in 755 agonists and 228 antagonists of the human *µ*OR. In cases where such description was unavailable, we used a labeling scheme based on the intrinsic activity value *E*_max_ determined by the [^35^S]GTP*γ*S functional assay which measures the activation level of G protein subsequent to the occupation of human *µ*OR by a chemical compound. This is achieved by measuring the binding of [^35^S]GTP*γ*S to the G_*α*_ subunit. ^16^ A compound was considered agonistic if *E*_max_ *>* 10% when the measurement was conducted in reference to DAMGO (i.e., 100% *E*_max_). When *E*_max_ was measured in reference to the basal activity (as 0%), we used *E*_max_ *>* 14% to label a compound as agonistic. *E*_max_ ≤ 14% was used to classify a compound as antagonistic. As an example, naltrexone’s *E*_max_ has been reported as approximately 8%^44^ or 14%^45^ in the literature. We manually curated the data and corrected any errors. Based on the above scheme, out of a total of 983 compounds, 755 and 228 were labeled as agonists and antagonists, respectively. The final data consists of molecular structures, represented by SMILES notation, and labels designating antagonists as the positive class and agonists as the negative class.

For tri-training, we collected 15,816 unique compounds that bind with the *µ, κ*, or *δ* opioid receptors in human, rat, or mouse from the open database BindingDB. ^38^ Compounds overlapping with the original labeled data set were removed. These compounds were used as unlabeled data for tri-training.

For external validation, we identified 15 agonists and 11 antagonists that are not in the training set, including from the work of Disney et al.,^46^ Pasternak and Pan,^24^ as well as the database NCATS Inxight drugs.^47^

### Protocol of building the tree-based classifiers

We performed data shuffling and stratified splitting of the data set into 90% for training/CV and 10% for unseen hold-out testing. PyCaret (version 2.3.10) ^31^ was used to build the random forest (RF), extra tree (ET) and XG boost (XGB) tree models for binary classification. For each compound, we first calculated 208 two-dimensional descriptors using the open-source package RDKit (version 2022.09.1).^23^ We then removed categorical features with statistically insignificant variances and dropped highly correlated numerical features (remove multicollinearity=true, multicollinearity threshold of 0.9). To avoid model overfitting, we tested different sized subsets of features (feature selection = true; n feature to select varied between 0.2 and 1). The latter parameter allowed the feature space to vary between 49 and 91. To address the class imbalance (agonist:antagonist ratio is about 3.3), an oversampling technique called SMOTE (Synthetic Minority Oversampling Technique)^32^ was employed. The hyperparameters were tuned to maximize the area under the curve (AUC) of the receiver operating characteristic (ROC) curve by performing a random grid search in 1000 iterations. The 9-fold stratified cross-validation was performed. The entire process was repeated 100 times for each feature selection threshold for statistical analysis. Once the best tree models were selected, they were retrained using the complete dataset for external validation.

### Protocol of building the MPNN classifiers

Chemprop (version 1.5.2) ^34^ was used to build the MPNNs for binary classification. The stratified data splitting was performed in the same manner as for the tree models. For each compound, a molecular graph was created based on the SMILES notation, and a feature vector was initialized based on atom and bond features computed by RDKit.^23^ The models were trained for 100 epochs with a batch size of 50. The loss function was binary cross-entropy, and the evaluation metric was F1. To prevent over-fitting, we used early stopping and performed 10-fold stratified cross-validation. To enhance model performance, we added 200 molecule-level features calculated and normalized by RDKit,^23^ generated an ensemble of classifiers trained on folds, and fine-tuned hyperparameters using 30 iterations of Bayesian optimization. Following hyperparameter tuning, we set the number of message-passing steps to 6, the neural network hidden size to 500, the number of feed-forward layers to 2, and the dropout probability to 0.1. We also optimized the classification probability threshold to maximizes the F1 score (Eq. 6). The resulting model was then tested on the 10% hold-out data. The above procedure (data splitting, training/CV, and test) was repeated 50 times for statistical analysis.

To interpret the predictions made by the MPNN classifiers, we applied the Monte Carlo Tree Search algorithm implemented in Chemprop^34^ and calculated the Tanimoto similarity scores using the topological fingerprints and MACCS keys with RDKit. ^23^ We also performed *k*-means clustering to identify representative substructures of the antagonistic compounds. For external validation, we trained Model 6 with the complete dataset.

### A tri-training scheme for binary classification

We employed PyCaret^31^ and Chemprop 1.5.2^34^ to build ET, XGB, and D-MPNN models using the same methodology as in the previous sections. We then predicted for the agonist/antagonist function on a new, unlabeled data set comprised of 15,816 unique ligands of the *µ, κ*, or *δ* opioid receptor found in the human, rat, and mouse experiments taken from bindingDB.^38^ The results of the classification for each of the three models (A, B, C) was used to construct new data sets for each model with the following protocol. A tri-training iteration begins by using the three models to make predictions on the unlabeled data set. For every unlabeled data point, if model A predicts a classification label that is different from the other two models (B and C), that data point with the label given by B and C will be added to model A’s training set. If all three models agree, the data point will be discarded. This prediction process is repeated until the unlabeled data set is exhausted, i.e., either moved to the training sets of the three models or discarded. Subsequently, all three models will be retrained with the inclusion of the new data points. The new models are evaluated on the unseen data. This completes the first iteration. The discarded data will be used for the next iteration of tri-training starting from the three new models. This procedure is repeated until all three models converge, i.e. any remaining data points in the unlabeled data set are given the same labels by all three models, or no significant improvement in F1 is observed. This disagreement scheme is included to reduce overfitting. This is known in the literature as tri-training with disagreement. In order to generate statistics, we performed three runs of tri-training and each run started with a different data splitting (90% for training/CV and 10% for hold-out testing).

### Model evaluation metrics

To evaluate model performance, we calculated the recall and precision for both classes (agonists and antagonists), which are defined as follows

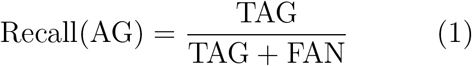

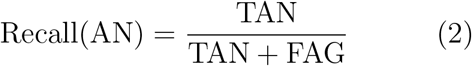

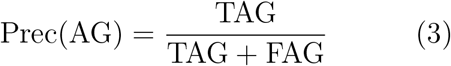

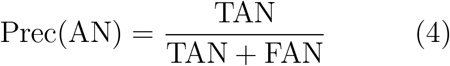

where TAG, TAN, FAG, and FAN denote the number of true agonists, true antagonists, false agonists, and false antagonists, respectively. Since the training data set is imbalanced, we labeled antagonists (the minority class) positives and agonists (the majority class) negatives to focus model training on antagonists. The overall performance of the classifier can be assessed by the balanced accuracy (BA), which is the arithmetic mean of the recalls for both classes.

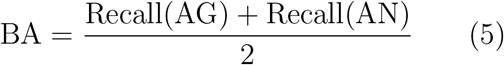

The F1 score, which is the harmonic mean of the recall and precision for the positives or the minority class (antagonists), was used to optimize the classification threshold.

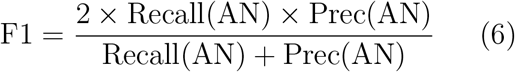

### Clustering analysis

The agglomerative clustering algorithm in Scikit-learn, with a Tanimoto distance (1 - Tanimoto similarity index) cutoff of 0.7 was used. The Tanimoto indices were calculated using ChemPro.^34^

## Supporting Information

A table of the evaluation metrics for the random forest and extra trees models. A table listing the 10 highest ranked features in the SHAP beeswarm plot of the ET model. A table summarizing the six successively optimized MPNNs. A table comparing the evaluation metrics for the initial and optimized MPNNs. A table comparing the true and predicted labels for the external dataset. Common structural motifs manually selected for structural classification of compounds in the training dataset. A figure of the cluster size vs. clustering label for the training dataset. A figure showing the probability distributions of the 10 highest ranked features in the SHAP plot. A figure showing the molecular structures of some morphine derivatives in the training dataset.

## Data Availability

The complete training and external validation datasets as well as models and tri-training data and Python scripts are freely downloadable at https://github.com/JanaShenLab/opioids. The training and external validation datasets can also be found at https://github.com/Myongin/muopioids.

## Acknowledgement

J.S. is supported by the National Institutes of Health (R35GM148261 and R01CA256557). Myongin Oh was supported by the ORISE fellowship. We thank Dr. Lei Shi and Dr. Michael Baumann from National Institute on Drug Abuse for the helpful discussion on the opioid activity measurements.

## Author Information

### Author Contributions

LS, JS, and MO designed the study. LS and MO collected the labeled data. MO carried out the tree-based and MPNN model training and wrote the initial draft manuscript. RL helped automate tree-based classification tasks and performed additional tests of the tri-training scheme. MS collected the unlabeled data, designed and executed the tri-training task, and prepared the corresponding manuscript content. All authors provided critical feedback on the data analysis. JS and LS revised the manuscript.

### Author disclaimer

This article reflects the views of the authors and should not be construed to represent FDA’s views or policies. The mention of commercial products, their sources, or their use in connection with material reported herein is not to be construed as either an actual or implied endorsement of such products by the Department of Health and Human Services.

## Notes

### Competing Interest Statement

The authors have declared no competing interest.

### Summary of Updates

We revised some discussion and removed 4 data points from the external validation set.

