## Supplementary material for "Machine Learned Classification of Ligand Intrinsic Activities at Human *µ*-Opioid Receptor": SI

Table S1: Summary of the evaluation metrics for the random forest and extra trees models with different feature selection thresholds

| | Random Forest | | | | Extra Trees | | | |
|---|---|---|---|---|---|---|---|---|
| $\alpha$ | $N$ | AUC (%) | BA (%) | F1 (%) | $N$ | AUC (%) | BA (%) | F1 (%) |
| 0.2 | 49 | 91.4±3.0 | 83.0±4.3 | 74.9±6.4 | 49 | 91.1±3.6 | 82.3±4.7 | 73.6±6.7 |
| 0.3 | 61 | 91.4±3.5 | 82.7±4.9 | 74.3±7.2 | 61 | 91.1±3.1 | 81.5±5.0 | 72.9±7.4 |
| 0.4 | 70 | 91.0±3.8 | 83.1±5.2 | 74.5±7.4 | 69 | 90.4±3.5 | 80.7±5.0 | 71.3±7.5 |
| 0.5 | 75 | 91.2±3.3 | 81.7±5.1 | 72.6±7.2 | 76 | 90.3±3.6 | 81.7±5.1 | 72.8±7.3 |
| 0.6 | 82 | 91.5±3.2 | 82.6±5.0 | 73.9±6.9 | 82 | 91.3±3.4 | 82.7±4.7 | 74.7±7.1 |
| 0.7 | 86 | 91.5±3.6 | 82.4±5.1 | 73.8±7.6 | 86 | 90.5±3.6 | 81.3±4.9 | 72.0±7.2 |
| 0.8 | 90 | 91.8±2.7 | 82.7±4.5 | 74.0±6.3 | 90 | 91.5±3.9 | 83.3±5.0 | 75.4±7.2 |
| 0.9 | 91 | 91.4±3.5 | 82.5±4.6 | 74.0±6.8 | 91 | 91.2±3.3 | 82.6±4.8 | 74.3±7.1 |
| 1.0 | 91 | 91.6±3.0 | 82.5±3.8 | 74.3±5.1 | 91 | 91.0±3.4 | 82.5±5.0 | 74.1±7.4 |

$\alpha$ denotes the feature selection threshold (parameter feature_selection_threshold in PyCaret[S1]). $N$ denotes the number of selected features. The average and standard deviations of the AUC-ROC, balanced accuracy (BA), and F1 score from the unseen tests are given.

Table S2: Description of the 10 highest ranked features in the SHAP beeswarm plot of the ET model ($\alpha = 0.8$)

| **Descriptor** | **Meaning** |
|---|---|
| $N_{\text{AlHc}}$ | Number of aliphatic heterocycles |
| $N_{\text{ArHc}}$ | Number of aromatic heterocycles |
| $N_{\text{XCCNR}}$ | Number of XCCNR groups |
| $N_{\text{ArRi}}$ | Number of aromatic rings |
| $N_{\text{AlOH}}$ | Number of aliphatic OH groups |
| $N_{\text{SaRi}}$ | Number of saturated rings |
| $N_{\text{imd}}$ | Number of imidazole groups |
| PEOE_VSA6 | MOE charge VSA descriptor 6 |
| MaxESI | Maximum electrotopological state index that combines electronic, topological, and valence states |
| SlogP_VSA8 | MOE logP VSA descriptor 8 |

Table S3: Summary of the six successively optimized MPNNs

| Model | 1 | 2 | 3 | 4 | 5 | 6 |
|---|---|---|---|---|---|---|
| Threshold optimization | × | ✓ | × | ✓ | ✓ | ✓ |
| Molecule-level features | × | × | ✓ | ✓ | ✓ | ✓ |
| Hyperparameter optimization | × | × | × | × | × | ✓ |
| $k$ | 10 | 10 | 10 | 10 | 5 | 5 |

Check mark indicates if a particular strategy, i.e., classification threshold optimization, addition of RDKit molecule-level features, or hyperparameter tuning is applied. $k$ is the number of classifiers in an ensemble.

Table S4: Summary of the evaluation metrics for the initial and successively optimized MPNNs

| **Model** | Recall(AN) | Recall(AG) | Prec(AN) | Prec(AG) | BA | F1(AN) |
|---|---|---|---|---|---|---|
| 1 | 71.7±10.0 | 94.5±2.5 | 80.5±7.7 | 91.7±2.7 | 83.1±5.1 | 75.4±7.3 |
| 2 | 73.4±9.8 | 93.1±3.0 | 77.2±7.6 | 92.1±2.7 | 83.3±4.9 | 74.8±6.7 |
| 3 | 72.4±10.5 | 94.9±2.4 | 81.7±7.1 | 92.0±2.9 | 83.7±5.3 | 76.4±7.4 |
| 4 | 75.5±10.7 | 93.5±3.3 | 78.8±8.3 | 92.6±2.9 | 84.5±5.3 | 76.6±7.4 |
| 5 | 75.1±10.5 | 93.5±3.3 | 78.7±8.4 | 92.5±2.9 | 84.3±5.3 | 76.4±7.4 |
| 6 | 76.3±10.6 | 93.9±3.1 | 80.2±8.2 | 93.0±2.9 | 85.1±5.0 | 77.5±6.7 |

The average and standard deviations of the metrics from 50 data splitting and training/CV runs are given in percentage. The models are explained in Table S3.

Table S5: Summary of the true and predicted labels for external dataset

| Compound | MPNN | ET | True Class |
|---|---|---|---|
| 6$\beta$-naltrexol | AN | AN | Antagonist |
| $\alpha$-naloxol | AN | AN | Antagonist |
| (17S)-methylnaltrexone | AN | AN | Antagonist |
| Naloxazone | AN | AN | Antagonist |
| Apigenin | AN | AN | Antagonist |
| 4',7-dihydroxyflavone | AN | AN | Antagonist |
| 4'-hydroxyflavanone | ~~AG~~ | ~~AG~~ | Antagonist |
| PF-4455242 | AN | ~~AG~~ | Antagonist |
| Bevenopran | AN | ~~AG~~ | Antagonist |
| Nalodeine | AN | ~~AG~~ | Antagonist |
| Orvinol 14 | ~~AG~~ | AN | Antagonist |
| Mitragynine | AG | AG | Agonist |
| Oxpheneridine | AG | AG | Agonist |
| Asimadoline | AG | AG | Agonist |
| Dermorphin | AG | AG | Agonist |
| 3-monoacetylmorphine | AG | AG | Agonist |
| Eseroline | AG | AG | Agonist |
| $(+/-)$-salsolinol | AG | AG | Agonist |
| Codeinone | AG | AG | Agonist |
| Phenazocine | AG | AG | Agonist |
| Morphinone | AG | AG | Agonist |
| Zilpaterol | AG | AG | Agonist |
| Morphiceptin | ~~AN~~ | ~~AN~~ | Agonist |
| Sophocarpidine | ~~AN~~ | ~~AN~~ | Agonist |
| Matrine | ~~AN~~ | ~~AN~~ | Agonist |
| Buprenorphine hemiadipate | AG | ~~AN~~ | Agonist |

AG and AN stand for agonist and antagonists, respectively. Incorrect predictions are indicated with cancel signs. The true labels (16 agonists in blue and 11 antagonists in red) are taken from the work of Disney et al.,[S2] Pasternak and Pan,[S3] and NCATS Inxight drugs.[S4]

Table S6: Summary of the AUCs of the hold-out test and external validation from three runs of tri-training with disagreement

| Iter | Data size | XGB (%) Test | XGB (%) Ext | Data size | ET (%) Test | ET (%) Ext | Data size | MPNN (%) Test | MPNN (%) Ext |
|---|---|---|---|---|---|---|---|---|---|
| Run 1 | | | | | | | | | |
| 0 | 886 | 90.6 | 69.6 | 886 | 86.4 | 80.9 | 886 | 89.7 | 83.0 |
| 1 | 1683 | 94.2 | 87.8 | 1673 | 93.1 | 70.9 | 5416 | 88.8 | 66.0 |
| 2 | 2287 | 89.5 | 81.2 | 2834 | 90.1 | 67.9 | 6173 | 89.4 | 68.5 |
| 3 | 2417 | 91.9 | 73.9 | 2861 | 91.4 | 74.2 | 6351 | 88.4 | 64.2 |
| 4 | 2455 | 91.5 | 79.1 | 2865 | 91.9 | 66.1 | 6489 | 90.4 | 68.5 |
| 5 | 2477 | 94.9 | 75.2 | 2872 | 92.3 | 67.9 | 7005 | 89.8 | 66.6 |
| Run 2 | | | | | | | | | |
| 0 | 886 | 84.4 | 72.7 | 886 | 90.5 | 75.2 | 886 | 92.3 | 83.0 |
| 1 | 1755 | 92.9 | 76.4 | 1727 | 90.1 | 53.9 | 6436 | 92.6 | 84.2 |
| 2 | 2977 | 80.7 | 61.2 | 2312 | 84.5 | 67.0 | 6796 | 90.6 | 76.4 |
| 3 | 2991 | 78.8 | 55.2 | 2344 | 84.6 | 67.3 | 6883 | 92.8 | 79.4 |
| 4 | 2995 | 88.6 | 57.6 | 2363 | 83.5 | 62.1 | 6886 | 93.1 | 80.6 |
| 5 | 2996 | 85.0 | 57.9 | 2373 | 82.7 | 72.4 | 6894 | 93.7 | 78.2 |
| Run 3 | | | | | | | | | |
| 0 | 886 | 86.3 | 61.8 | 886 | 82.3 | 81.5 | 886 | 93.3 | 80.6 |
| 1 | 2263 | 87.0 | 81.2 | 1705 | 85.1 | 56.1 | 4592 | 94.5 | 69.1 |
| 2 | 2681 | 93.7 | 78.8 | 3387 | 85.3 | 82.1 | 4979 | 95.4 | 75.2 |
| 3 | 2737 | 94.4 | 79.4 | 3396 | 84.6 | 76.4 | 5181 | 95.2 | 72.1 |
| 4 | 2771 | 88.4 | 64.2 | 3397 | 84.0 | 76.1 | 5283 | 95.7 | 72.1 |
| 5 | 2784 | 93.5 | 70.9 | 3398 | 83.4 | 72.8 | 5391 | 95.1 | 72.7 |

Iteration number is given in the first column, where iteration 0 refers to the initial models. The initial XGB and ET were trained and cross-validated using 90% of the dataset, while MPNN was taken from the models 02, 33, and 41 of the previous training (Table 2 in the main text). Column Data size refers to the total number of data points in the training/CV; among them 886 are labeled and the rest are pseudo-labeled data. Column Test and Ext refer to the hold-out test (10% of the dataset) and the external validation using 26 compounds in Table S5.

### Supplemental figures

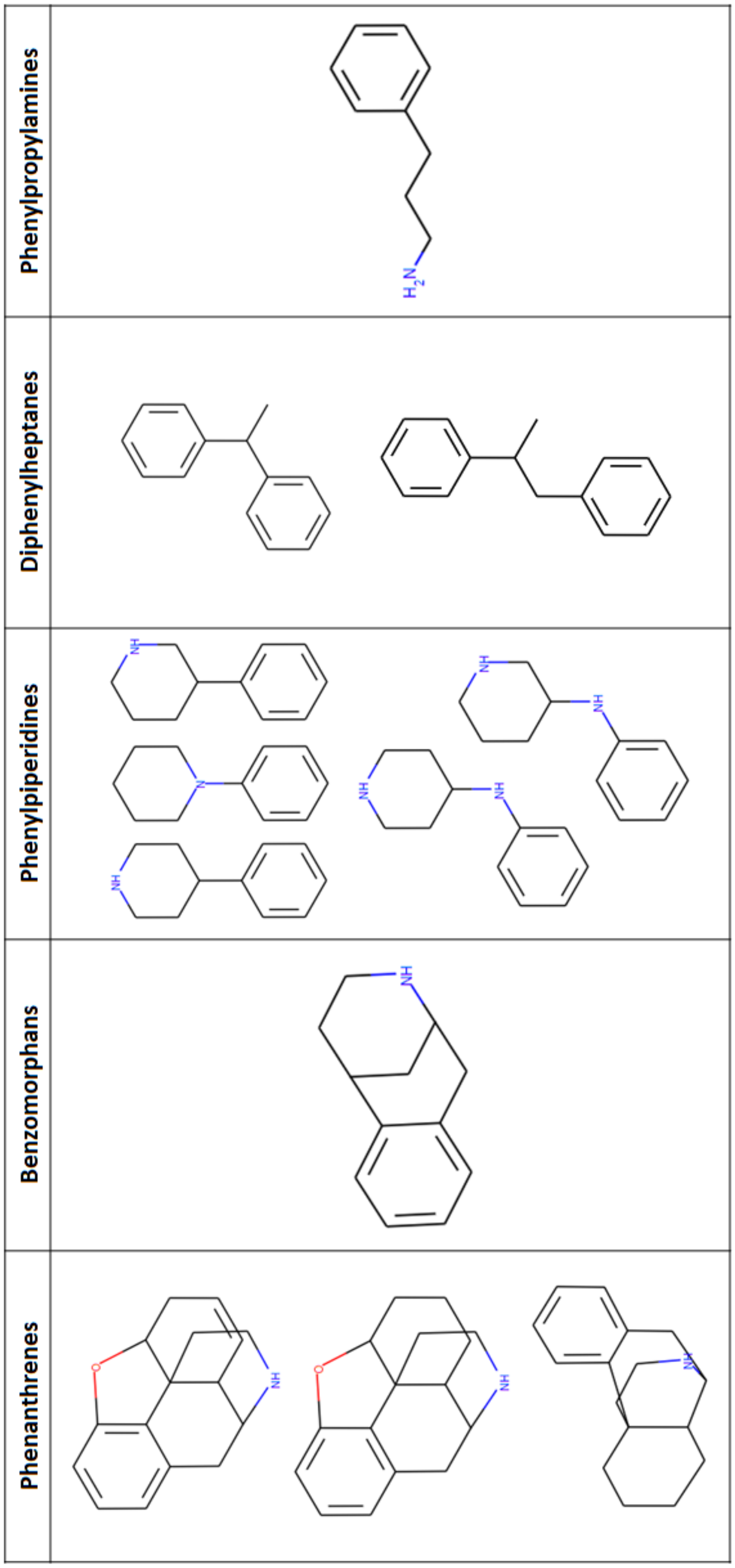


Figure S1: Common structural motifs manually selected for structural classification of compounds in the training dataset

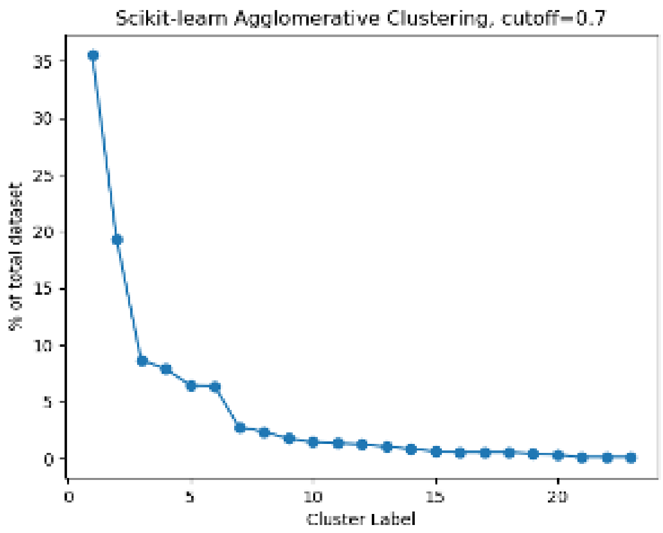


Figure S2: Cluster size vs. clustering label for the training dataset. The clustering analysis was performed on the SMILE strings of the training dataset molecules using the agglomerative method in Scikit-Learn. The clustering metric was the Tanimoto distance (1-Tanimoto similarity) of 0.7 between two molecules.

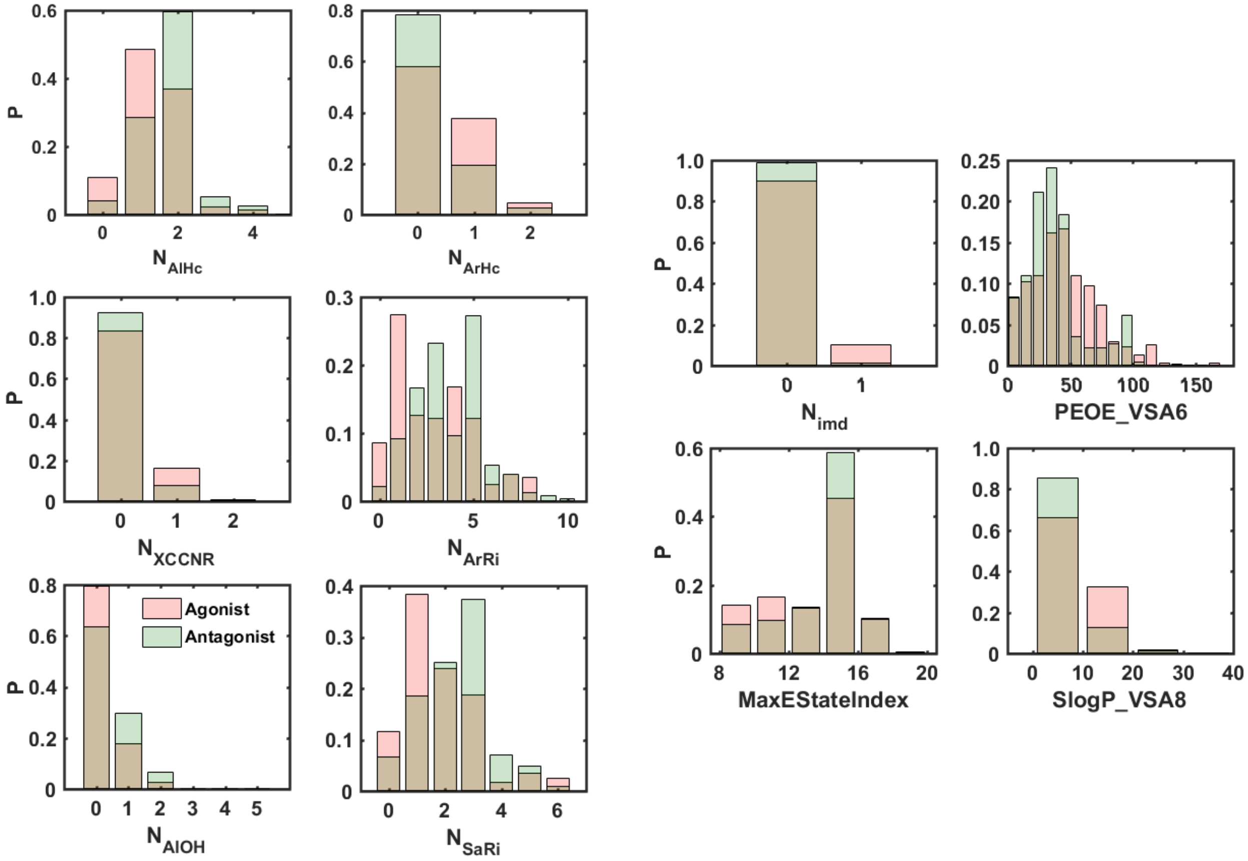


Figure S3: Normalized probability distributions of the 10 highest ranked features in the SHAP beeswarm plot of the best performing ET model for agonists (red) and antagonists (green)

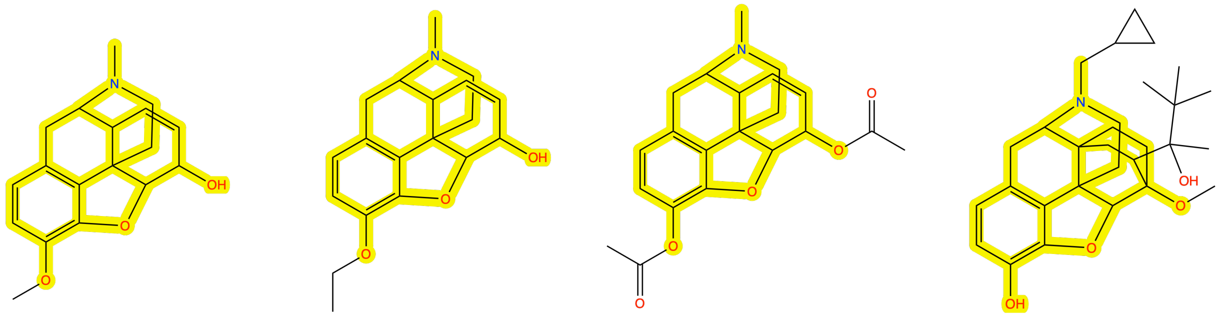

Methylmorphine (codeine) Ethylmorphine Diacetylmorphine (heroin) Buprenorphine

Figure S4: Molecular structures of some morphine derivatives in the training dataset. The morphine substructure is highlighted.

### References


(S1) Ali, M. PyCaret: An Open Source, Low-Code Machine Learning Library in Python. 2020.

(S2) Disney, A.; Olson, K. M.; Shafer, A. M.; Moore, S. C.; Anand, J. P.; Traynor, J. R.; Husbands, S. M. Opioid Antagonists from the Orvinol Series as Potential Reversal Agents for Opioid Overdose. *ACS Chem. Neurosci.* **2022**, *13*, 3108–3117.

(S3) Pasternak, G. W.; Pan, Y.-X. Mu Opioids and Their Receptors: Evolution of a Concept. *Pharmacol. Rev.* **2013**, *65*, 1257–1317.

(S4) Siramshetty, V. B.; Grishagin, I.; Nguyen, D.-T.; Peryea, T.; Skovpen, Y.; Stroganov, O.; Katzel, D.; Sheils, T.; Jadhav, A.; Mathé, E. A.; Southall, N. T. NCATS Inxight Drugs: A Comprehensive and Curated Portal for Translational Research. *Nucleic Acids Res.* **2022**, *50*, D1307–D1316.
